# Robust machine learning for *Clostridioides difficile* typing from mass spectrometry via data augmentation

**DOI:** 10.1101/2024.10.29.620907

**Authors:** Alejandro Guerrero-López, Mario Blázquez-Sánchez, Lucía Bravo-Antón, Lucía Schmidt-Santiago, Carlos Sevilla-Salcedo, David Rodríguez-Temporal, Belén Rodríguez-Sánchez, Vanessa Gómez-Verdejo

## Abstract

Machine learning (ML) approaches applied to Matrix-Assisted Laser Desorption Ionization–Time of Flight Mass Spectrometry (MALDI-TOF MS) spectra have shown promise for the typing of *Clostridioides difficile*, yet their deployment in routine clinical settings remains challenging due to strong sensitivity to acquisition variability. Differences in culture media, incubation time, protein extraction protocols, and instrumentation across hospitals often lead to substantial performance degradation when models are evaluated under heterogeneous or previously unseen conditions.

In this work, we systematically analyze the impact of methodological and technical variability on ML-based *C. difficile* typing and investigate whether data augmentation (DA) strategies can mitigate these effects. Using a dedicated dataset of 60 isolates acquired under diverse conditions, we show that DA substantially improves robustness to variability when training on spectra from selective *C. difficile* agar media. Importantly, models trained with DA achieve performance levels approaching those obtained using enriched Schaedler agar media, while relying exclusively on standard 24-hour incubation.

Evaluation on an independent cohort of 28 newly acquired isolates confirms that DA significantly reduces performance degradation under real-world domain shift. To facilitate adoption and reproducibility, we release MALDIDA, an open-source Python library for DA of MALDI-TOF MS spectra.

## 1. Introduction

Matrix-Assisted Laser Desorption Ionization–Time of Flight Mass Spectrometry (MALDI-TOF MS) has revolutionized clinical microbiology in the last decade. This technique allows for rapid and accurate bacterial identification by analyzing their protein profiles within minutes [1], unlike traditional methods, which can take several days and require highly trained personnel in molecular biology methods, including Pulsed Field Gel Electrophoresis (PFGE) and Polymerase Chain Reaction (PCR) Ribotyping [2]. The analysis of MALDI-TOF MS spectra typically relies on attributes such as peak height and position, focusing primarily on ribosomal proteins that are species-specific biomarkers [3, 4]. Nevertheless, for more complex tasks such as antibiotic resistance detection or strain-level discrimination, visual analysis is insufficient [5, 6]. Consequently, automatic classification of MALDI-TOF MS spectra using machine learning (ML) models has gained increasing attention as a cost-effective and scalable alternative for bacterial characterization [2, 7, 8, 9].

In this study, MALDI-TOF MS and ML methods were evaluated for the automatic differentiation of *Clostridioides difficile* ribotypes (RTs). *C. difficile*, a Gram-positive, anaerobic, spore-forming species transmitted through the fecal-oral route, is the leading cause of antibiotic-associated infectious diarrhea in hospitals [10, 11]. Certain RTs are associated with increased pathogenicity and hospital outbreaks [12, 13], such as RT027 [14, 15], considered hypervirulent due to toxin hyperproduction [11]. Other hypervirulent RTs, including RT181, have emerged in recent years and are increasingly reported across Europe [16].

One of the main challenges in automatically classifying clinical samples using MALDI-TOF MS spectra is the limited availability of data [7]. In addition, MALDI-TOF MS is prone to substantial variability, including inconsistencies in the presence or absence of specific peaks and fluctuations in their intensities [17, 18]. This variability is further exacerbated by differences in growth conditions, protein extraction (PE) protocols, laser intensity, machine calibration, and operator expertise. Such lack of reproducibility is a well-recognized limitation in bacterial characterization, affecting the reliability and generalization of ML models across laboratories [19, 20, 18].

While enriched media (e.g., Schaedler agar) tend to produce more robust and consistent spectral data, their use is impractical for routine clinical workflows due to additional isolation and incubation steps. In contrast, selective media (e.g., CHROMID® selective agar), although essential for isolating *C. difficile* from fecal samples [21, 22, 23, 24], introduce increased spectral variability and reduced ML classification performance.

As a result, ML models trained on MALDI-TOF MS spectra tend to be biased towards the acquisition characteristics of the training domain, leading to performance degradation when evaluated under heterogeneous or previously unseen conditions—a phenomenon commonly referred to as domain shift [25, 26, 27]. Ensuring robustness and generalization under such conditions is therefore a critical requirement for the reliable deployment of ML systems in clinical practice.

From a ML perspective, this work systematically investigates acquisition-induced domain shift in MALDI-TOF MS-based classification and evaluates data augmentation (DA) as a strategy to improve robustness and generalization. We designed a dedicated dataset comprising 60 *C. difficile* isolates cultivated under diverse methodological and technical conditions to explicitly capture both methodological variability (culture media and PE protocols) and technical variability (instrumentation and acquisition time). Using this dataset, we conducted a two-fold study: (i) we analyzed how specific sources of variability affect the generalization performance of commonly used ML models; and (ii) we evaluated whether domain-aware DA strategies can mitigate performance degradation when training data are limited or acquired under heterogeneous protocols.

To this end, we implemented a DA pipeline designed to model realistic spectral perturbations, including peak shifting and new peak generation, acting as a regularization mechanism against domain shift. We demonstrate that DA primarily improves robustness and performance stability rather than average performance. Crucially, evaluation on an independent held-out cohort of 28 newly acquired isolates confirms that DA significantly mitigates performance degradation under realistic deployment conditions. To facilitate reproducibility and adoption, we release MALDIDA, an open-source Python library for DA of MALDI-TOF MS data. A graphical abstract summarizing the study objectives and methodology is presented in Figure 1.

**Figure 1.**
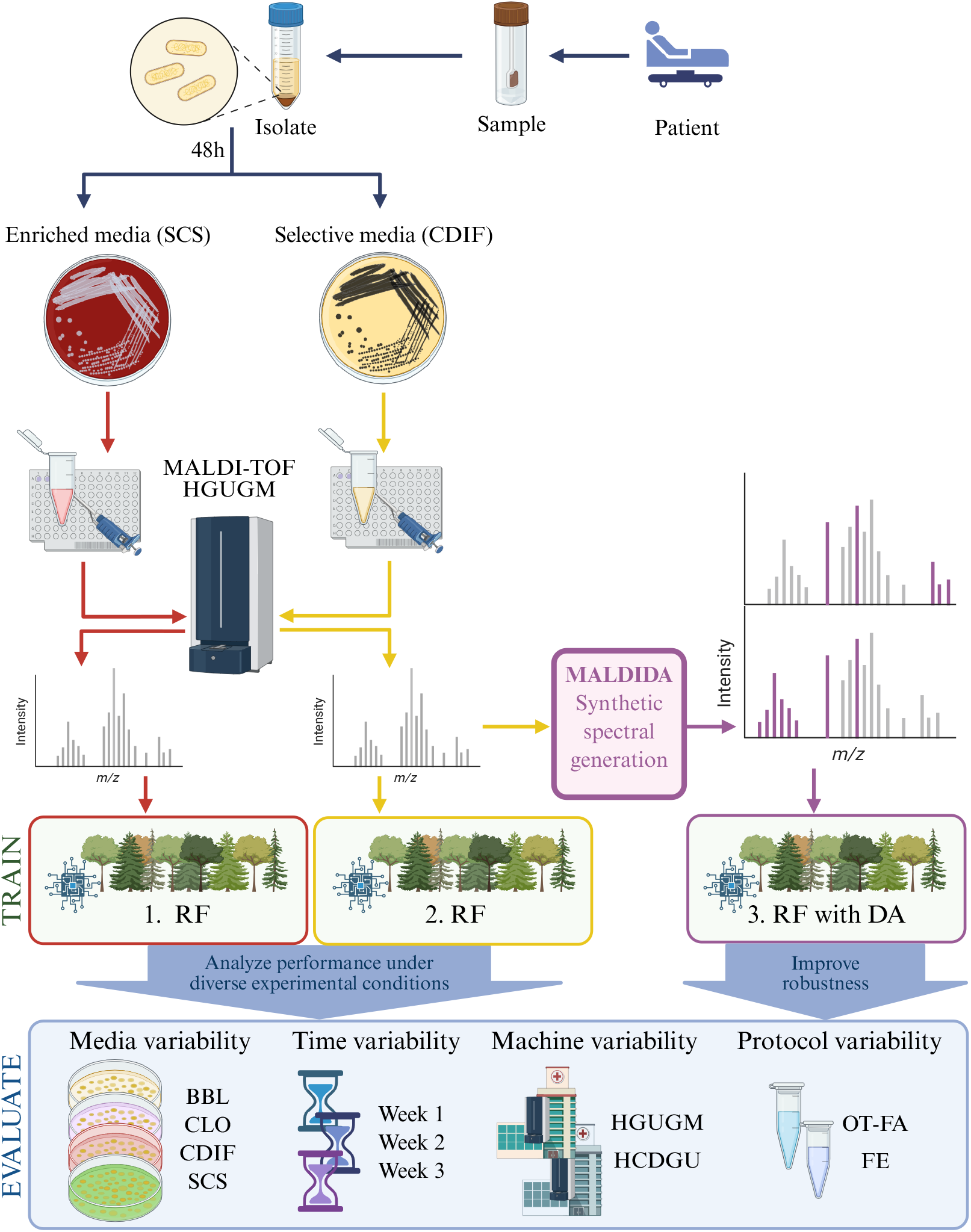
Overall workflow. Samples of six *C. difficile* RTs were initially cultured on CHROMID® selective agar (CDIF) and enriched Schaedler agar (SCS) at 37ºC anaerobically for 48 h. Mass spectra were then obtained via MALDI-TOF MS from both media, without protein extraction. Three classification scenarios were trained: (1) a Random Forest (RF) model on SCS spectra, (2) an RF model on CDIF spectra, and (3) an RF model on CDIF spectra augmented with our proposed data augmentation (DA) techniques. Each model was subsequently evaluated for robustness against various sources of variability, including culture media, intra-hospital variability, instrumentation, and protein extraction protocol.

## 2. Methods

### 2.1. MALDI-TOF MS spectra

To assess how different conditions affect MALDI-TOF MS spectra, we analyzed 60 *C. difficile* clinical isolates retrieved from an institutional strain repository. The isolates were originally recovered from stool specimens of patients with diarrhea across 10 Spanish hospitals; the corresponding specimens were tested using the Xpert® *C. difficile* BT assay, and RTs were assigned by PCR ribotyping [28, 29, 30]. The collection comprised 10 isolates each of ribotypes RT023, RT027, RT078, RT106, RT165, and RT181. For each isolate, MALDI-TOF MS spectra were repeatedly extracted under varying laboratory conditions, including intrahospital variability extracted in three different weeks (Time), types of culture medium (Media), PE protocols, and mass spectrometers used in different hospitals (Hosp). Moreover, for inter-spot reproducibility, each isolate was analyzed in three different spots (triplicates) following the same procedure. Hence, we have 180 mass spectra (3 replicates per each one of the 60 isolates) for each condition analyzed in this study. In all scenarios, all isolates were incubated anaerobically at 37 °C for 48 hours. Spectra were acquired using 1 µL of organic HCCA (*α*-Cyano-4-hydroxycinnamic acid) matrix, in the positive ion mode within the 2,000 to 20,000 Da range.

For time and media variability, the samples were evaluated over three weeks using the on-target formic acid extraction protocol (OT-FA) on four culture media: CHROMID® *C. difficile* selective agar (CDIF) from bioMérieux, enriched Schaedler agar (SCS), enriched *Brucella* agar (BBL) from Becton Dickinson, and selective *Clostridium* agar (CLO) from bioMérieux. The use of the OT-FA protocol was also compared with the full extraction protocol (FE) on the same medium for the selective CDIF media.

All these analyses were conducted exclusively at the Hospital General Universitario Gregorio Marañón (HGUGM) using MALDI-TOF MS technology with an MBT Smart MALDI Biotyper (Bruker Daltonics, Bremen, Germany). To test hospital variabilities, all isolates were processed using the OT-FA extraction protocol in CDIF at another hospital, Hospital Central de la Defensa Gómez Ulla (HCDGU), performed with its MALDI Biotyper® sirius System (Bruker Daltonics), that is, different from HGUGM, but with the same MALDI-TOF MS technology.

Subsequently, to assess the robustness of our approach under real deployment conditions, we included a second, independent set of 28 clinical isolates, balanced across the six RTs, for independent evaluation of the ML models. For this cohort, incubation and processing conditions were randomized to emulate a realistic routine scenario. Concretely, for each isolate we randomly assigned (i) one of three culture media: BBL, CLO, and the new media Columbia agar (COS) from bioMérieux; and (ii) one of two PE protocols (OT-FA or FE). The three technical replicates acquired per isolate were generated under the same assigned medium–protocol combination to preserve within-isolate reproducibility, while ensuring heterogeneity across isolates. Spectra were acquired in a MALDI Biotyper® sirius system (Bruker Daltonics®) at HCDGU in a different acquisition time point.

### 2.2. Preprocessing and binning

Each MALDI-TOF MS spectrum is represented by a set of *k* tuples of mass-to-charge ratio and intensity:

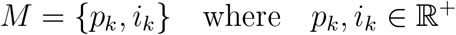

where *p*_*k*_ denotes the position of the *k*-th peak (in *m/z*, measured in Daltons) and *i*_*k*_ denotes its corresponding intensity, both constrained to positive real numbers. For the preprocessing, we started applying the conventional open-source software MALDIquant [31] to preprocess MALDI-TOF MS, but using the version in Python code extracted from [32], which consists of five steps: (1) a intensity transformation with a square-root method to stabilize the variance, (2) a smoothing using the Savitzky-Golay algorithm with half-window-size of 10, (3) a baseline removal which is estimated with 10 iterations of the SNIP algorithm, (4) an intensity calibration using the total ion current (TIC), and (5) a spectra trimming to values in the 2,000–20,000 Da m/z range. Thus, each MALDI-TOF MS is a set of non-discrete tuples (*p*_*k*_), (*i*_*k*_) in an 18,000 Da range, where *k* is variable. Figure 2 shows the result of all these steps.

**Figure 2.**
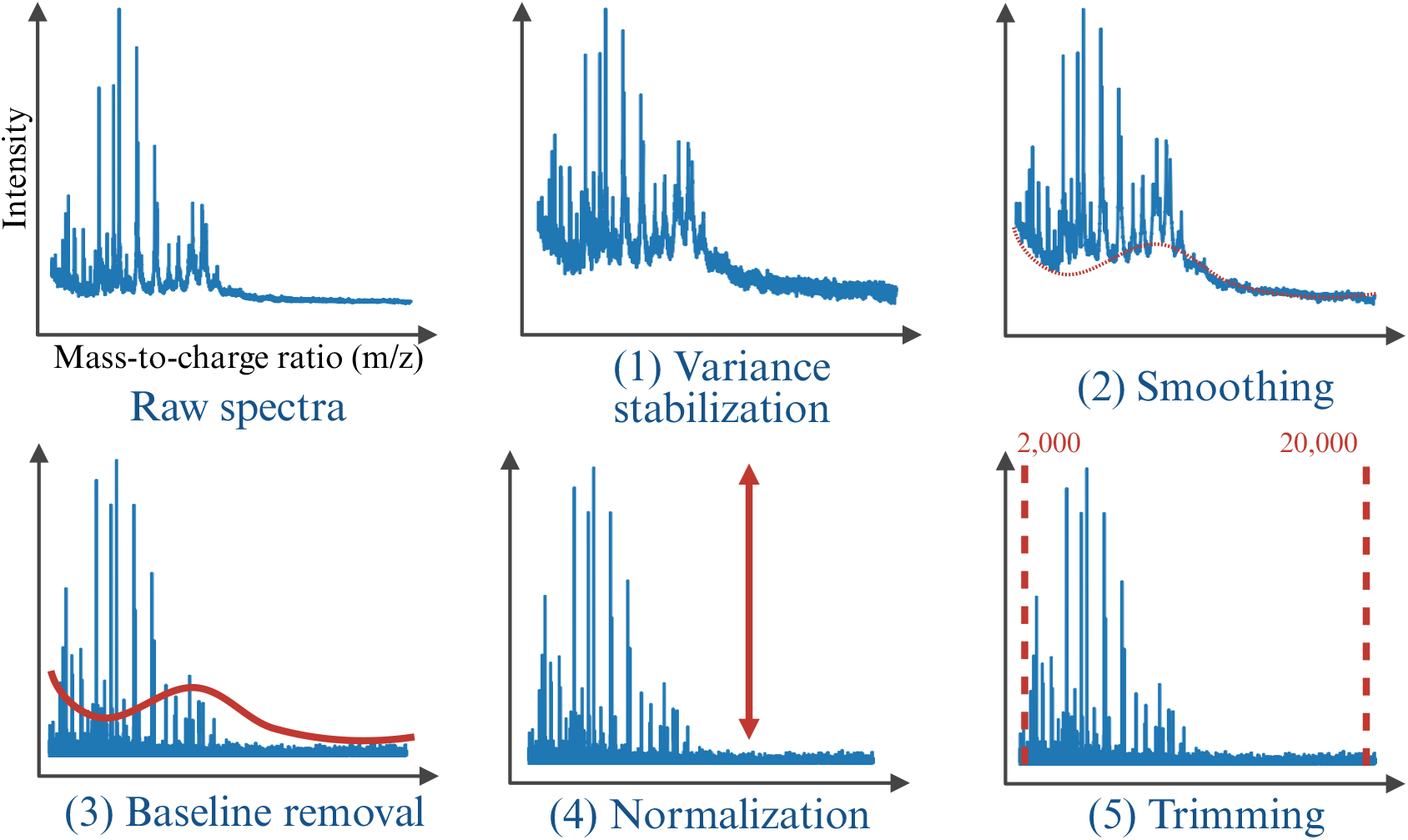
Summary of the preprocessing steps.

After preprocessing, following previous works [2, 33], we constructed the feature vectors of fixed-length required by ML methods by applying a binning step. To perform the binning, the m/z axis was partitioned in the range of 2,000 to 20,000 m/z into equal-sized bins of 3 m/z size, and the intensity of all measurements in the sample falling into the same bin was summed up. Thus, each sample was represented by a fixed-dimensionality vector containing 6,000 features.

### 2.3. Data augmentation

The analysis of MALDI-TOF MS spectra differences across media, time, PE protocols, and machine acquisition for the same sample reveals significant spectral variability, as shown in Figure 3 for an RT027 sample. This includes the presence of new peaks (as shown for Week 2 and Week 3 in CDIF medium in Fig. 3c), changes in intensity (as shown when testing on different hospital machines in Fig. 3a), and shifts in positions (as shown for BBL media and Week 2 in Fig. 3b). Such variability can lead to ML models biased to the acquisition conditions, resulting in poor generalization when faced with acquisition differences in the test set. To improve the robustness of these methods and given the small sample size in which this variability may not be fully represented, we propose simulating these variations through DA.

**Figure 3.**
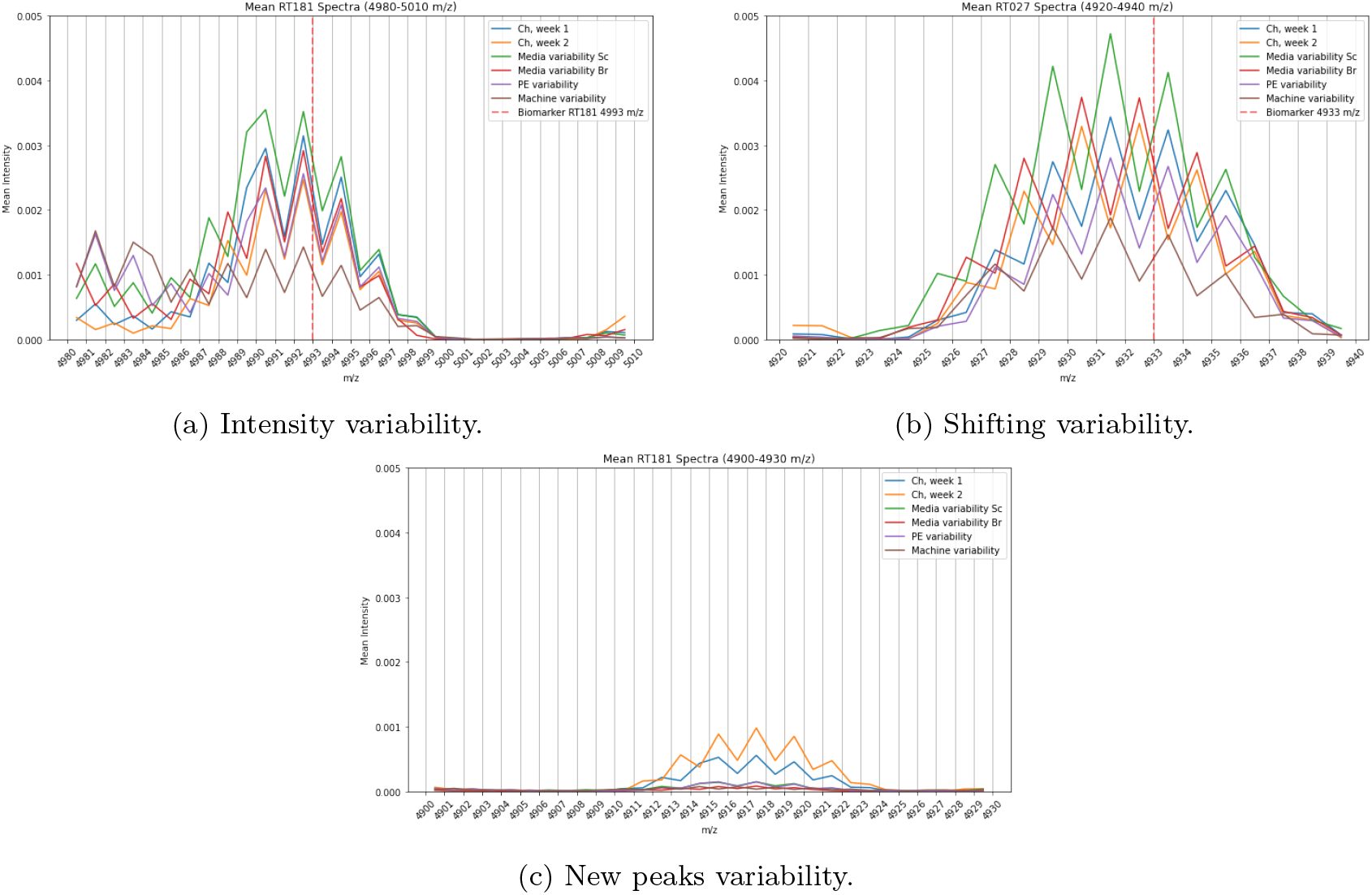
Differences in spectra of the same ribotype (RT) due to variability conditions: a) the mean of all RT181 samples where each variable studied offer a different value of intensity peak; b) the mean of all RT027 samples where a well-known biomarker 4,933 m/z [34, 35] is shifted 2-3 Da depending on the variable condition; c) mean of all RT181 samples where a new peak appears only on selective CDIF media.

In particular, we propose to generate synthetic spectra that can reproduce these variations by modifying training data spectra with three techniques:

1. **Intensity variations**: To simulate the variability in intensities observed between different machines and weeks, we perturb a random subset of the peaks. For the *k*-th peak (*p*_*k*_) with a positive intensity *i*_*k*_, we perturb it with probability *p*_1_. Formally, let

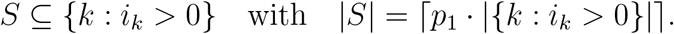

Then, the augmented intensity 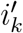 is defined as

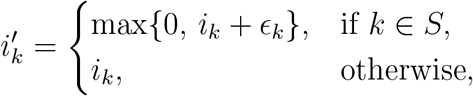

where *ϵ*_*k*_ ∼𝒩 (0, *v*_1_) is Gaussian noise with zero mean and variance *v*_1_. In this way, random noise is added to a fraction *p*_1_ of the peaks while ensuring that the intensities remain non-negative following the behaviour of Fig. 3a.

**2. Peak shifting**: To simulate variability due to instrumental conditions, a random fraction *p*_2_ of peaks is selected for shifting. Formally, let

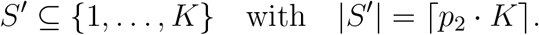

For each peak *k*, the new position is defined as

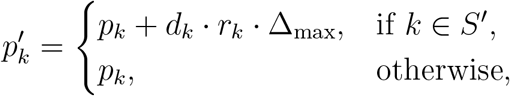

where *d*_*k*_ ∈ {−1, +1} is a random sign, *r*_*k*_ is the normalized position of the peak (e.g.,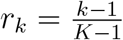), and

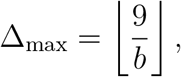

with *b* being the bin size (so that the maximum shift is up to ±9 Da). Finally, the intensities *i*_*k*_ are re-interpolated on the original *p*_*k*_ grid to yield the augmented intensities following the behaviour of Fig. 3b.

**3. New peaks**: To simulate machine variability, we perturb a random fraction *p*_3_ of the zero-intensity locations. Let

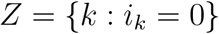

and choose a subset

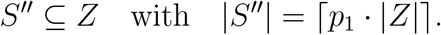

Then, for each *k* ∈ *S*^′′^, the augmented intensity is defined as

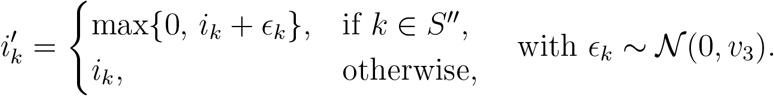

This procedure introduces new peaks by adding low-variance Gaussian noise to regions with zero intensity, following the behaviour of Fig. 3c.

### 2.4. Experimental setup

To study the effect of MALDI-TOF MS variability on different classifier methods, we designed an experiment aimed at classifying each isolate into one of six *C. difficile* RTs (RT181, RT165, RT106, RT078, RT027, and RT023), and we involved several state-of-the-art ML models commonly used in the field [36]: (i) a Support Vector Machine (SVM) with the most common kernels, such as the linear and Radial Basis Function (RBF) kernels; (ii) K-Nearest Neighbors (KNN); (iii) a Random Forest (RF); and (iv) a Light Gradient Boosting Machine (LGBM). The hyperparameters of these models were adjusted using a 5-fold cross-validation technique. For the SVM, hyperparameter C was cross-validated between 0.01 and 100, and kernel width (*γ*), when using RBF, was cross-validated covering a logarithmic range from to 1,000. The number of estimators of the RF was evaluated within a range of 50 to 300. Furthermore, a range of 3 to 20 neighbour values was cross-validated for KNN.

In routine microbiology laboratories, when cultures are performed, *C. difficile* is commonly recovered on selective media (e.g., CDIF or CLO) to facilitate isolation from stool specimens. In line with the hospital’s standard diagnostic procedures, our baseline model (referred to as Baseline CDIF) was trained on Week 1 data acquired with selective medium CDIF, OT-FA protocol, and processed on the HGUGM machine. In contrast, a second model (referred to as Baseline SCS) mirrors the Baseline CDIF model, but uses enriched medium SCS. Although enriched media are designed to promote optimal bacterial growth, they require an initial culture on a selective medium to isolate candidate bacteria, resulting in an additional time of 48 hours before acquiring MALDI-TOF MS spectra. Therefore, in this first experiment, we evaluate how variability affects both media.

We evaluated each model’s Area Under the Curve (AUC) performance for the prediction of RT under different variability conditions using an outer Leave-One-Isolate-Out (LOIO) strategy. In this approach, the model was trained on all isolates from a chosen baseline domain (either Baseline CDIF or Baseline SCS), while a single isolate was held out for testing. Importantly, all technical replicates of the held-out isolate were selected from one of several distinct conditions, allowing us to assess how well the model generalizes on methodological and technical variability. These conditions were:

- Baseline condition: The test isolate came from the same domain as the training data (Baseline CDIF or Baseline SCS), representing the ideal scenario.
- Media variability: The test isolate was chosen from a different medium than the training data, specifically from one of three alternatives (BBL, CLO, or SCS/CDIF), ensuring it was always distinct from the baseline.
- PE variability: The test isolate varied in terms of the use of OT-FA versus FE protocol.
- Time variability: The test isolate was selected from a different week (Week 2 or Week 3), to evaluate the model’s robustness to temporal changes.
- Machine variability: The test isolate was sourced from a different hospital and generated using a different MALDI-TOF MS machine.

It is important to note that the test isolate was never included in the training data and was solely used for testing purposes. This LOIO process was repeated for each isolate in the data set, ensuring that each isolate was used as a test case in turn.

## 3. Results

### 3.1. Variability analysis regarding model performance

Tables 1 and 2 present the AUC values for the Baseline CDIF and Baseline SCS models, respectively. In each table, the first row shows the baseline AUC obtained by testing the models on samples acquired under the same conditions as the training data (culture medium, PE method, same week, and machine). The subsequent rows report the AUC when one condition is varied at a time. Each column corresponds to a specific model, with the final column providing the mean AUC across models, and the last row summarizes the mean variability for each model.

**Table 1:**
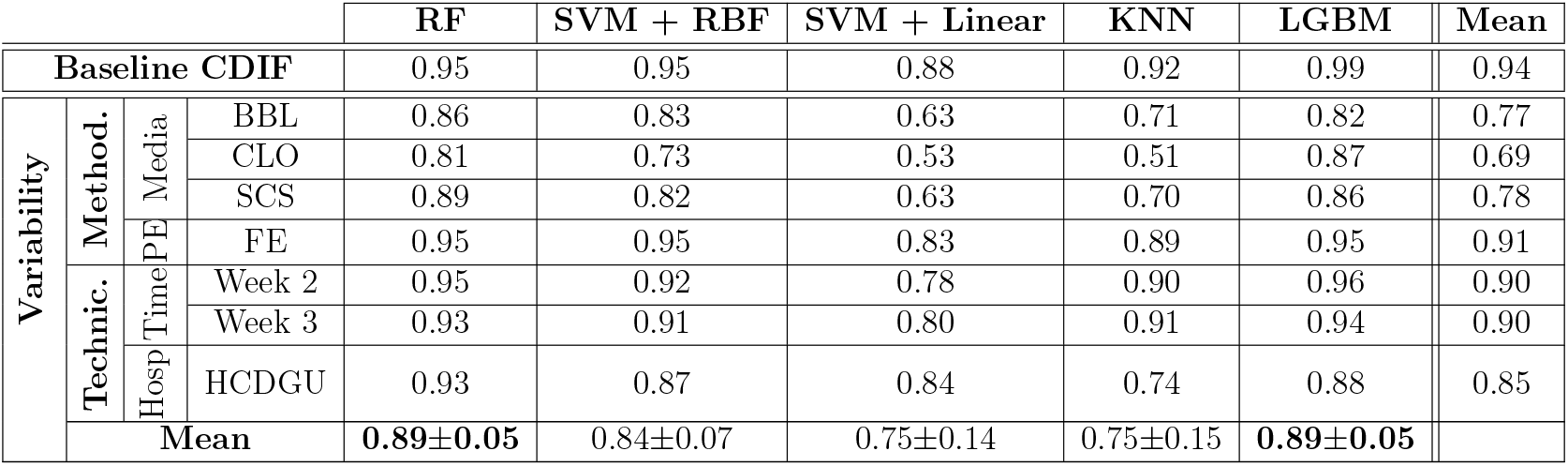
Area Under the Curve (AUC) scores comparing the performance of the Baseline CDIF model against various methodological and technical variations across different models: Random Forest (RF), Support Vector Machine (SVM) with different kernels, K-Nearest Neighbors (KNN), and Light Gradient Boosting Machine (LGBM). The Baseline CDIF model was trained using Week 1 data, selective CDIF medium, the OT-FA protocol, and the HGUGM instrument. Results in bold indicate the classifier that performs best overall across all variability scenarios.

**Table 2:** Area Under the Curve (AUC) scores comparing the performance of the Baseline SCS model against various methodological and technical variations across different models: Random Forest (RF), Support Vector Machine (SVM) with different kernels, K-Nearest Neighbors (KNN), and Light Gradient Boosting Machine (LGBM). The Baseline SCS model was trained using Week 1 data, enriched SCS medium, the OT-FA protocol, and the HGUGM instrument. Results in bold indicate the classifier that performs best overall across all variability scenarios.

|  |  |  |  | RF | SVM + RBF | SVM + Linear | KNN | LGBM | Mean |
| --- | --- | --- | --- | --- | --- | --- | --- | --- | --- |
| Baseline SCS |  |  |  | 0.99 | 0.97 | 0.78 | 0.91 | 0.98 | 0.93 |
| Variability | Meth. | Media | BBL | 0.97 | 0.93 | 0.70 | 0.84 | 0.98 | 0.88 |
|  |  |  | CLO | 0.94 | 0.89 | 0.56 | 0.68 | 0.84 | 0.78 |
|  |  |  | CDIF | 0.94 | 0.81 | 0.63 | 0.67 | 0.84 | 0.78 |
|  | Tec. | Time | Week 2 | 0.97 | 0.95 | 0.69 | 0.86 | 0.95 | 0.88 |
|  |  |  | Week 3 | 0.96 | 0.93 | 0.69 | 0.82 | 0.91 | 0.86 |
|  | Mean |  |  | 0.96±0.02 | 0.90±0.07 | 0.66±0.08 | 0.77±0.12 | 0.90±0.07 |  |

In Table 1 (using the CDIF medium), changing the sample acquisition time led to an average AUC reduction of 4 points for Weeks 2 and 3. Altering the culture medium resulted in a decrease of up to 25 AUC points when using the selective medium CLO. Variability due to different hospital equipment caused a 9-point reduction in AUC. The LGBM model had a baseline AUC of 0.99, which decreased to 0.89 when variability was introduced.

Table 2 (using the enriched SCS medium) shows that the RF model’s baseline AUC decreased from 0.99 to 0.96 under average variability conditions, while the LGBM model decreased to an AUC of 0.90. Note that the use of enriched media in a real hospital setting requires prior isolation of *C. difficile* on selective media (CDIF), adding an extra 24–48 hours to the total growth time.

Thus, these findings confirm the lack of robustness of current ML models to the technical and methodological variations commonly encountered in clinical practice, thereby limiting their widespread application in microbiology laboratories and routine diagnostics.

### 3.2. Evaluation of the DA techniques

In this section, we conducted an ablation study to analyze the impact of each DA step (see Section 2.3) on the RF classifier’s performance. The objective of this analysis was to identify the optimal configuration for the DA process by systematically evaluating the contribution of each transformation. For this experiment, the number of generated DA samples was varied from 100 to 2,500. The following strategies and parameter ranges were evaluated:

- **Intensity variations:** Different values of *p*_1_ (25%, 50%, 75%) and noise standard deviations *v*_1_ (0.01, 0.05, 0.1, 0.5, 1) were evaluated.
- **Peak shifting:** Proportional shifting was applied using *p*_2_ values of 0.2, 0.4, 0.6, 0.8, and 1.
- **New peaks:** With a fixed *p*_3_ = 0.15, standard deviations *v*_3_ of 0.01, 0.05, 0.1, and 0.2 were analyzed.

The models were trained using the Baseline CDIF dataset and evaluated with an LOIO scheme. For each left-out isolate, the model was tested on the remaining mass spectra (across four media, two extraction protocols, and two hospital machines, excluding the one used in training).

Figure 4 compares the performance of the RF model with the different DA strategies against the baseline RF model without DA. Intensity variability modification resulted in a reduction of overall AUC (Fig. 4a). In contrast, shifting peak positions produced consistent increases in AUC as the number of augmented samples increased (Fig. 4b). Similarly, the addition of new peaks in zero-intensity regions improved performance. Finally, a combined evaluation, where the number of augmented samples was increased from 0 to 6,000 (approximately 100 times the original dataset of 60), showed that performance improvements plateaued after roughly 3,000 additional samples (Fig. 4d). Therefore, it appears suitable that combining peak position shifting and the addition of zero-intensity peaks, with a target augmentation size of approximately 3,000 samples, provides an effective configuration for the DA process.

**Figure 4.**
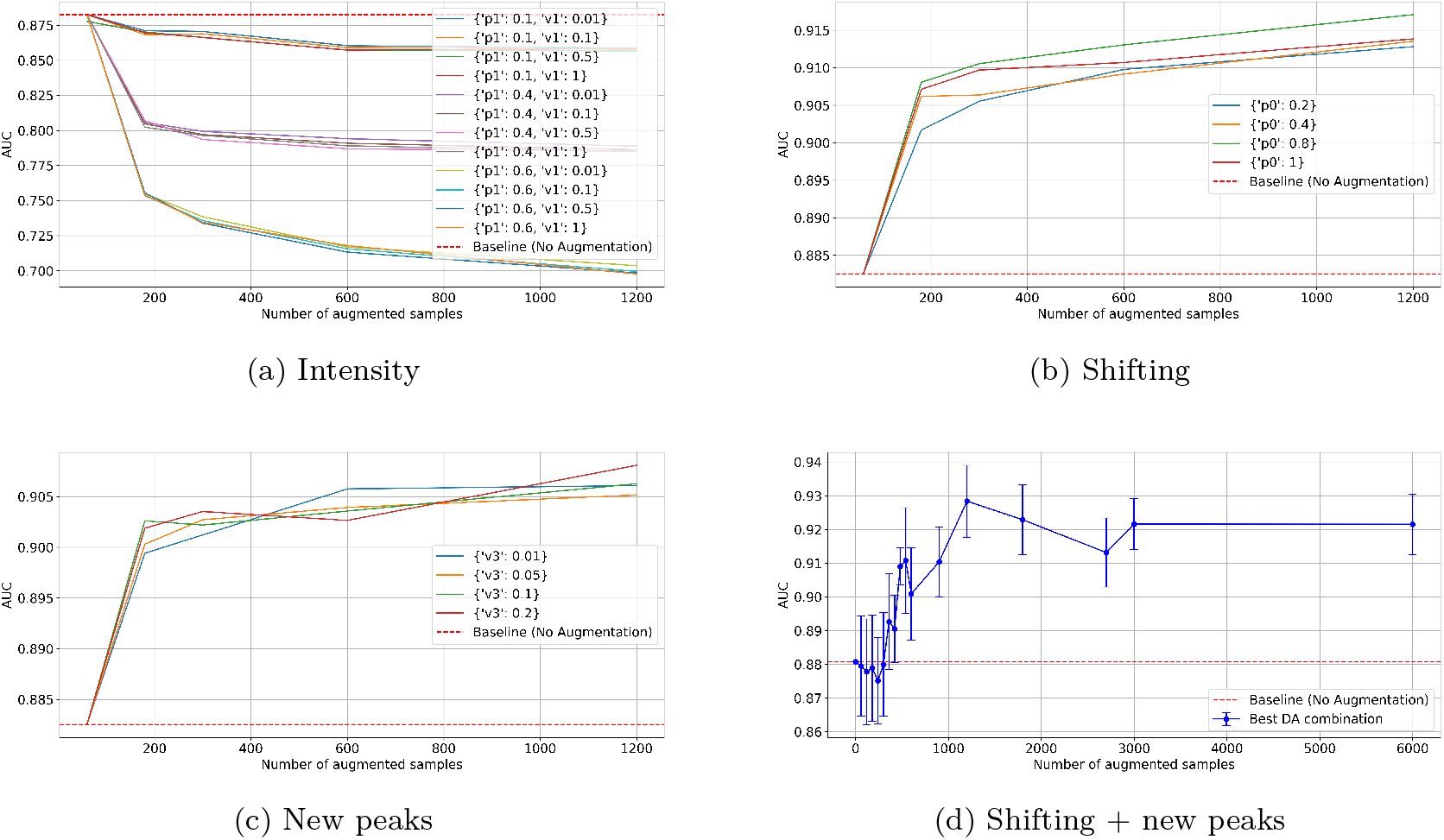
Ablation study for the data augmentation (DA) techniques. Subfigures (a-c) analyze the Area Under the Curve (AUC) of a random forest (RF) trained with the different DA techniques proposed, while subfigure d) shows the AUC scores of the RF trained by combining the two best DA techniques.

To facilitate reproducibility and adoption, we provide these DA strategies through MALDIDA, an open-source Python library designed for targeted data augmentation of MALDI-TOF MS spectra. The library is modular and user-friendly, allowing practitioners to easily customize augmentation steps (e.g., peak shifting and new peak generation) and tune parameters according to their specific acquisition conditions and downstream tasks.

### 3.3. ML model robustness with DA

In this section, we first investigated whether applying DA techniques to the selective CDIF medium can limit the performance decline of ML models due to variability. Based on the enriched SCS medium results presented in Table 2, we aimed to obtain comparable results by training a classifier with samples acquired with the medium (faster acquisition) combined with DA synthetic samples. For this evaluation, we focus on the RF model, which showed the most consistent performance across variability conditions in Section 3.1, and tested its performance with respect to the AUC.

Table 3 compares three configurations: the Baseline CDIF RF model trained without DA, the same model trained with 3,000 augmented samples generated using shifting and new peak techniques, and a reference model trained on spectra acquired from enriched SCS medium without DA.

**Table 3:** Comparison of Area Under the Curve (AUC) scores for the Baseline CDIF random forest (RF) trained with data augmentation (DA) (w/ DA) versus without DA (w/o DA), evaluated under different sources of methodological and technical variability. Results obtained using enriched SCS medium w/o DA are shown as a reference. The Baseline CDIF reference uses fixed conditions: Week 1, selective CDIF medium, OT-FA protocol, and HGUGM machine. Results in bold indicate the configuration that performs best overall across all variability scenarios.

|  |  |  |  | Baseline CDIF | Baseline CDIF | Baseline SCS |
| --- | --- | --- | --- | --- | --- | --- |
|  |  |  |  | w/ DA (RF) | w/o DA (RF) | w/o DA (RF) |
| Baseline |  |  |  | **0.96** | 0.95 | 0.99 |
| Variability | Meth. | Media | BBL | 0.89 | 0.86 | 0.97 |
|  |  |  | CLO | 0.88 | 0.81 | 0.94 |
|  |  |  | SCS | 0.91 | 0.89 | 0.94 |
|  | Technic. | Time | Week 2 | 0.98 | 0.92 | 0.97 |
|  |  |  | Week 3 | 0.98 | 0.93 | 0.96 |
|  |  | Hosp | HCDGU | 0.93 | 0.93 | - |
| | Mean | | | **0.93 ± 0.04** | 0.89 ± 0.05 | 0.96 ± 0.02 |

Regarding temporal variability, the application of DA increased the AUC from 0.92 to 0.98 in both Weeks 2 and 3. Concerning methodological variability across culture media, the baseline RF model achieved AUC values of 0.86, 0.81, and 0.89 for BBL, CLO, and SCS media, respectively; with DA, these values increased to 0.89, 0.88, and 0.91. Finally, for hospital-related technical variability, DA did not affect performance, with both configurations yielding an AUC of 0.93.

Importantly, the CDIF model trained with DA achieves mean AUC values (0.93 ± 0.04) that are comparable to, and in some conditions approach, those obtained using the enriched SCS medium (0.96 ± 0.02), despite relying exclusively on spectra acquired after standard 24 h incubation on selective media.

Secondly, to evaluate the robustness of our approach under real deployment conditions, we trained the classifiers exclusively on the original CDIF training cohort and conducted an independent evaluation over a collection of 28 new isolates. As detailed in Section 2.1, this new cohort was balanced across the six RTs and subjected to randomized incubation and processing conditions to emulate a realistic routine scenario and ensure technical and methodological heterogeneity.

For this experiment, the independent cohort was used solely as an external test set, without any form of fine-tuning, calibration, or domain adaptation using target-domain data. Predictions were generated at the spectrum level using the three technical replicates. This evaluation protocol ensures that all reported results reflect true out-of-distribution generalization, closely mimicking a realistic deployment scenario in which new samples are processed under unseen laboratory conditions.

Table 4 summarizes the isolate-level performance obtained on an independent test cohort acquired under heterogeneous and previously unseen conditions. As expected, a marked degradation in performance is observed when models trained under controlled settings are evaluated on this external dataset, highlighting the severity of domain shift in real-world deployments. Under these challenging conditions, training with augmented spectra by the proposed DA pipeline consistently improves robustness across acquisition settings.

**Table 4:** Isolate-level classification Area Under the Curve (AUC) using random forest on the independent test cohort. Models were trained exclusively on the original CDIF dataset from Hospital General Universitario Gregorio Marañón (HGUGM) with OT-FA. Results are reported with (w/) and without (w/o) data augmentation (DA), across culture media (BBL, CLO, COS) and protein extraction protocols (OT-FA, FE), for both hospital machines (HGUGM and Hospital Central de la Defensa Gómez Ulla (HCDGU)).

|  | HGUGM |  |  |  |  |  | HCDGU |  |  |  |  |  |
| --- | --- | --- | --- | --- | --- | --- | --- | --- | --- | --- | --- | --- |
|  | FE |  |  | OT-FA |  |  | FE |  |  | OT-FA |  |  |
|  | BBL | CLO | COS | BBL | CLO | COS | BBL | CLO | COS | BBL | CLO | COS |
| **w/ DA** | 1.00 | 0.75 | 0.82 | 1.00 | 0.85 | 1.00 | 1.00 | 0.75 | 0.78 | 0.97 | 1.00 | 0.83 |
| **w/o DA** | 0.96 | 0.78 | 0.80 | 1.00 | 0.70 | 0.92 | 0.92 | 0.75 | 0.78 | 0.91 | 0.62 | 0.79 |

Overall, the systematic gains observed when using DA indicate that synthetic data generation constitutes an effective strategy to partially mitigate domain shift and improve model robustness in realistic, heterogeneous clinical environments, thereby enhancing real-world deployability.

## 4. Discussion

The design of a dedicated dataset comprising 60 isolates cultivated under diverse conditions, specifically engineered to capture technical variability (introduced by differences across instruments and acquisition time points) and methodological variability (resulting from the use of multiple culture media and PE protocols), has enabled a rigorous analysis of the significant impact of experimental variability on the performance of ML models for rapid *C. difficile* typing using MALDI-TOF MS spectra.

Our initial variability analysis revealed that while shifts in sample acquisition time and differences in hospital equipment lead to moderate declines in AUC, changes in the culture medium—which commonly vary across hospitals in routine clinical practice [19, 20, 18]—result in a pronounced performance drop of up to 25 AUC points. Both the RF and LGBM models, despite their high baseline performance under controlled conditions, showed considerable sensitivity to variations in culture protocols. In contrast, the enriched SCS medium consistently yielded higher AUC values; however, the requirement of an initial isolation step on selective media limits its practicality for routine use [23]. Importantly, although high classification performance can be achieved under controlled acquisition settings, our results reveal a substantial degradation when models are evaluated on independent cohorts acquired under heterogeneous and previously unseen conditions, highlighting the challenges associated with real-world deployment.

The ablation study of DA techniques further exemplified the challenges of replicating natural spectral variability. Specifically, modifying peak intensities, although widely used in SMOTE-based methods [37, 38], is a generic, non-targeted procedure that may reduce performance. In contrast, applying peak shifting consistently increased AUC, and the introduction of new peaks improved the model’s generalization capacity, both novel techniques proposed in this study. Notably, performance improvements plateaued beyond 3,000 augmented samples, indicating an optimal level of augmentation beyond which additional synthetic data yields no further improvement.

In a subsequent experiment, DA techniques were applied to spectra acquired from the selective CDIF medium to assess their ability to mitigate the adverse effects of acquisition variability. Using 3,000 augmented samples generated through peak shifting and new peak generation, the model exhibited substantial robustness gains. In particular, the AUC increased from 0.92 to 0.98 for temporal variability and from 0.95 to 0.98 for PE-related variability. Despite these improvements, variability associated with the culture medium itself remained more challenging, as performance did not fully reach the levels observed with the enriched SCS medium, and no significant gains were observed for equipment-induced variability. Overall, rather than simply increasing average performance, DA primarily acts as a regularization mechanism that reduces sensitivity to methodological and technical sources of heterogeneity, thereby improving model stability across acquisition conditions.

Finally, results on an independent test dataset highlight that domain-aware augmentation can substantially improve robustness when deploying MALDI-TOF MS classifiers across heterogeneous acquisition settings. The performance degradation observed in the baseline model confirms that spectral shifts induced by instrumentation and workflow variability pose a significant challenge for supervised models trained on data from a single domain. However, the gains obtained with synthetic spectra suggest that mimicking realistic perturbations helps the model internalize invariances that generalize beyond the training distribution. Importantly, the improvements were not uniform across classes, indicating that some RTs benefit more from augmentation than others, likely due to differences in spectral distinctiveness or sensitivity to acquisition variability. Notably, while DA does not uniformly improve performance across all conditions, it consistently enhances model robustness and generalization capability under heterogeneous acquisition settings. Overall, these findings support the idea that targeted, physics-guided DA can act as an effective and low-cost strategy to enhance the deployment reliability of MALDI-TOF MS–based ML systems in real clinical environments.

Beyond methodological robustness, our findings have direct practical implications for clinical microbiology workflows. In particular, the combination of selective CDIF media with DA enables performance levels that approach those obtained with enriched SCS media, while relying exclusively on spectra acquired after standard 24 h incubation. This suggests that DA can partially compensate for the increased variability associated with faster, routine culture protocols, reducing the need for longer incubation times without substantially sacrificing classification performance. As a result, the proposed approach represents a pragmatic trade-off between robustness, turnaround time, and deployability in real-world clinical settings.

From an ML perspective, these findings highlight that robustness to acquisition-induced domain shift, rather than average performance under controlled validation settings, should be considered a primary design and evaluation criterion for applied ML systems in clinical environments.

Our open-source MALDIDA Python library provides a practical toolset for implementing these augmentation techniques. Importantly, MALDIDA is designed to be flexible and can be extended to other spectrometry and spectroscopy domains that share similar peak-based representations, such as FTIR spectroscopy, where spectra are defined over wavenumbers and are affected by baseline drift and peak shifts due to acquisition conditions, or Raman spectroscopy, which similarly exhibits peak shifts and intensity variability across instruments and measurement setups. Future work will focus on optimizing DA parameters and exploring additional strategies to enhance the generalizability of diagnostic models across diverse clinical settings.

## Data availability

The reproducibility code is available at GitHub, and we make the DA techniques available as a Python package named MALDIDA, installed via pip from here. All data used in this study are available in the GitHub repository.

## References

[1] A. Tran, K. Alby, A. Kerr, M. Jones, P. H. Gilligan, Cost savings realized by implementation of routine microbiological identification by matrix-assisted laser desorption ionization–time of flight mass spectrometry, J. of clinical microbiology 53 (2015) 2473–2479.

[2] C. Weis, A. Cuénod, B. Rieck, O. Dubuis, S. Graf, C. Lang, M. Oberle, M. Brackmann, K. K. Søgaard, M. Osthoff, K. Borgwardt, A. Egli, Direct antimicrobial resistance prediction from clinical MALDI-TOF mass spectra using machine learning, Nature medicine 28 (2022) 164–174.

[3] R. J. Pais, F. Sharara, R. Zmuidinaite, S. Butler, S. Keshavarz, R. Iles, Bioinformatic identification of euploid and aneuploid embryo secretome signatures in IVF culture media based on MALDI-ToF mass spectrometry, J. of Assisted Reproduction and Genetics 37 (2020) 2189–2198.

[4] T. Brignoli, M. Recker, W. W. Lee, T. Dong, R. Bhamber, M. Albur, P. Williams, A. W. Dowsey, R. C. Massey, Diagnostic MALDI-TOF MS can differentiate between high and low toxic Staphylococcus aureus bacteraemia isolates as a predictor of patient outcome, Microbiology 168 (2022) 001223.

[5] Q. Mao, X. Zhang, Z. Xu, Y. Xiao, Y. Song, F. Xu, Identification of Escherichia coli strains using MALDI-TOF MS combined with long short-term memory neural networks, Aging 16 (2024).

[6] T.-H. Lin, H.-Y. Chung, M.-J. Jian, C.-K. Chang, H.-H. Lin, C.-M. Yu, C.-L. Perng, F.-Y. Chang, C.-W. Chen, H.-S. Shang, Innovative strategies against superbugs: Developing an AI-CDSS for precise Stenotrophomonas maltophilia treatment, J. of Global Antimicrobial Resistance 38 (2024) 173–180.

[7] C. Weis, C. Jutzeler, K. Borgwardt, Machine learning for microbial identification and antimicrobial susceptibility testing on MALDI-TOF mass spectra: a systematic review, Clinical Microbiology and Infection 26 (2020) 1310–1317.

[8] L. Schmidt-Santiago, A. Guerrero-López, C. Sevilla-Salcedo, D. Rodríguez-Temporal, B. Rodríguez-Sánchez, V. Gómez-Verdejo, Machine learning applied to maldi-tof data in a clinical setting: a systematic review, bioRxiv (2025) 2025–01.

[9] M. Blázquez-Sánchez, A. Guerrero-López, A. Candela, A. Belenguer-Llorens, J. M. Moreno, C. Sevilla-Salcedo, M. Sánchez-Cueto, M. J. Arroyo, M. Gutiérrez-Pareja, V. Gómez-Verdejo, P. M. Olmos, L. Mancera, P. Muñoz, M. Marín, L. Alcalá, D. Rodríguez-Temporal, B. Rodríguez-Sánchez, Automated web-based typing of Clostridioides difficile ribotypes via MALDI-TOF MS, BMC Bioinformatics 26 (2025). doi:10.1186/s12859-025-06200-6.

[10] J. Czepiel, M. Dróżdż, H. Pituch, E. J. Kuijper, W. Perucki, A. Mielimonka, S. Goldman, D. Wultańska, A. Garlicki, G. Biesiada, Clostridium difficile infection, European J. of Clinical Microbiology & Infectious Diseases 38 (2019) 1211–1221.

[11] R. Markovska, G. Dimitrov, R. Gergova, L. Boyanova, Clostridioides difficile, a New “Superbug”, Microorganisms 11 (2023) 845.

[12] K. A. Davies, H. Ashwin, C. M. Longshaw, D. A. Burns, G. L. Davis, M. H. Wilcox, E. S. Group, et al., Diversity of Clostridium difficile PCR ribotypes in Europe: results from the European, multicentre, prospective, biannual, point-prevalence study of Clostridium difficile infection in hospitalised patients with diarrhoea (EUCLID), 2012 and 2013, Eurosurveillance 21 (2016) 30294.

[13] A. Gomez-Simmonds, M. K. Annavajhala, M. P. Nunez, N. Macesic, et al., Intestinal dysbiosis and risk of posttransplant clostridioides difficile infection in a longitudinal cohort of liver transplant recipients, mSphere 7 (2022) e00361–22. doi:10.1128/msphere.00361-22.

[14] M. He, F. Miyajima, P. Roberts, L. Ellison, D. J. Pickard, M. J. Martin, T. R. Connor, S. R. Harris, D. Fairley, K. B. Bamford, et al., Emergence and global spread of epidemic healthcare-associated Clostridium difficile, Nature genetics 45 (2013) 109–113.

[15] V. F. Viprey, G. L. Davis, A. D. Benson, D. Ewin, W. Spittal, J. J. Vernon, M. Rupnik, A. Banz, F. Allantaz, P. Cleuziat, et al., A point-prevalence study on community and inpatient Clostridioides difficile infections (CDI): results from Combatting Bacterial Resistance in Europe CDI (COMBACTE-CDI), July to November 2018, Eurosurveillance 27 (2022) 2100704.

[16] M. Kachrimanidou, S. Metallidis, O. Tsachouridou, C. Harmanus, V. Lola, E. Protonotariou, L. Skoura, E. Kuijper, Predominance of Clostridioides difficile PCR ribotype 181 in northern Greece, 2016–2019, Anaerobe 76 (2022) 102601.

[17] M. T. Olson, P. S. Blank, D. L. Sackett, A. L. Yergey, Evaluating reproducibility and similarity of mass and intensity data in complex spectra—applications to tubulin, J. of the American Society for Mass Spectrometry 19 (2011) 367–374.

[18] R. A. Giebel, C. R. Worden, S. Rust, G. T. Kleinheinz, M. Robbins, T. Sandrin, Microbial fingerprinting using matrix-assisted laser desorption ionization time-of-flight mass spectrometry (MALDI-TOF MS) applications and challenges., Advances in applied microbiology 71 (2010) 149–84.

[19] A. J. Saenz, C. E. Petersen, N. B. Valentine, S. L. Gantt, K. H. Jarman, M. T. Kingsley, K. L. Wahl, Reproducibility of matrix-assisted laser desorption/ionization time-of-flight mass spectrometry for replicate bacterial culture analysis, Rapid communications in mass spectrometry 13 (1999) 1580–1585.

[20] Olson, Matthew T and Epstein, Jonathan A and Sackett, Dan L and Yergey, Alfred L, Production of reliable maldi spectra with quality threshold clustering of replicates, J. of the American Society for Mass Spectrometry 22 (2011) 969–975.

[21] S. Luk, W. K. To, T. K. Ng, W. T. Hui, W. K. Lee, F. Lau, A. M. W. Ching, A cost-effective approach for detection of toxigenic Clostridium difficile: toxigenic culture using ChromID Clostridium difficile agar, J. of Clinical Microbiology 52 (2014) 671–673.

[22] J. J. Yang, Y. S. Nam, M. J. Kim, S. Y. Cho, E. You, Y. S. Soh, H. J. Lee, Evaluation of a chromogenic culture medium for the detection of Clostridium difficile, Yonsei medical journal 55 (2014) 994–998.

[23] J. H. Chen, V. C. Cheng, O.-Y. Wong, S. C. Wong, S. Y. So, W.-C. Yam, K.-Y. Yuen, The importance of matrix-assisted laser desorption ionization–time of flight mass spectrometry for correct identification of Clostridium difficile isolated from chromID C. difficile chromogenic agar, J. of Microbiology, Immunology and Infection 50 (2017) 723–726.

[24] M. M. Nerandzic, C. J. Donskey, Effective and reduced-cost modified selective medium for isolation of Clostridium difficile, J. of clinical microbiology 47 (2009) 397–400.

[25] A. Cesaro, S. C. Hoffman, P. Das, C. de la Fuente-Nunez, Challenges and applications of artificial intelligence in infectious diseases and antimicrobial resistance, npj Antimicrobials and Resistance 3 (2025) 2.

[26] Y. P. H. Chin, W. Song, C. E. Lien, C. H. Yoon, W.-C. Wang, J. Liu, P. A. Nguyen, Y. T. Feng, L. Zhou, Y. C. J. Li, D. W. Bates, Assessing the international transferability of a machine learning model for detecting medication error in the general internal medicine clinic: Multi-center preliminary validation study, JMIR Medical Informatics 9 (2021) e23454. doi:10.2196/23454.

[27] F. Tranchellini, Y. Farag, C. Jutzeler, L. Meegahapola, Evaluating deep learning sepsis prediction models in icus under distribution shift: A multi-centre retrospective cohort study, medRxiv (2025). doi:10.1101/2025.07.31.25332542, preprint.

[28] S. L. J. Stubbs, J. S. Brazier, G. L. O’Neill, B. I. Duerden, Pcr targeted to the 16s-23s rrna gene intergenic spacer region of¡i¿clostridium difficile¡/i¿ and construction of a library consisting of 116 different pcr ribotypes, Journal of Clinical Microbiology 37 (1999) 461–463. doi:10.1128/jcm.37.2.461-463.1999.

[29] A. Indra, S. Huhulescu, M. Schneeweis, P. Hasenberger, S. Kernbichler, A. Fiedler, G. Wewalka, F. Allerberger, E. J. Kuijper, Characterization of clostridium difficile isolates using capillary gel electrophoresis-based pcr ribotyping, Journal of Medical Microbiology 57 (2008) 1377–1382. doi:10.1099/jmm.0.47714-0.

[30] M. Marín, A. Martín, L. Alcalá, E. Cercenado, C. Iglesias, E. Reigadas, E. Bouza, Clostridium difficile isolates with high linezolid mics harbor the multiresistance gene ¡i¿cfr¡/i¿, Antimicrobial Agents and Chemotherapy 59 (2015) 586–589. doi:10.1128/aac.04082-14.

[31] S. Gibb, K. Strimmer, MALDIquant: a versatile R package for the analysis of mass spectrometry data, Bioinformatics 28 (2012) 2270–2271.

[32] G. De Waele, G. Menschaert, W. Waegeman, An antimicrobial drug recommender system using maldi-tof ms and dual-branch neural networks, Elife 13 (2024) RP93242.

[33] C. Weis, M. Horn, B. Rieck, A. Cuénod, A. Egli, K. Borgwardt, Topological and kernel-based microbial phenotype prediction from MALDI-TOF mass spectra, Bioinformatics (Oxford, England) 36 (2020) i30–i38.

[34] J. Corver, J. Sen, B. Hornung, B. Mertens, E. Berssenbrugge, C. Harmanus, I. Sanders, N. Kumar, T. Lawley, E. Kuijper, et al., Identification and validation of two peptide markers for the recognition of Clostridioides difficile MLST-1 and MLST-11 by MALDI-MS, Clinical Microbiology and Infection 25 (2019) 904–e1.

[35] A. Guerrero-López, C. Sevilla-Salcedo, A. Candela, M. Hernández-García, E. Cercenado, P. M. Olmos, R. Cantón, P. Muñoz, V. Gómez-Verdejo, R. Del Campo, et al., Automatic antibiotic resistance prediction in klebsiella pneumoniae based on maldi-tof mass spectra, Engineering Applications of Artificial Intelligence 118 (2023) 105644.

[36] X. A. López-Cortés, J. M. Manríquez-Troncoso, J. Kandalaft-Letelier, S. Cuadros-Orellana, Machine learning and matrix-assisted laser desorption/ionization time-of-flight mass spectra for antimicrobial resistance prediction: A systematic review of recent advancements and future development, J. of Chromatography A (2024) 465262.

[37] C. A. Astudillo, X. A. López-Cortés, E. Ocque, J. M. Manríquez-Troncoso, Multi-label classification to predict antibiotic resistance from raw clinical MALDI-TOF mass spectrometry data, Scientific Reports 14 (2024) 31283.

[38] V. Macaya Mejias, D. Zabala-Blanco, X. A. López-Cortés, F. Tirado, J. M. Manríquez-Troncoso, R. Ahumada-García, Predicting bacterial antibiotic resistance using MALDI-TOF mass spectrometry databases with ELM applications, J. of Computer Science & Technology 24 (2024).

